# Super-resolution microscopy reveals distinct epigenetic states regulated by estrogen receptor activity

**DOI:** 10.1101/2025.06.16.659976

**Authors:** Tara Akhshi, Shengen Shawn Hu, Esme Wheeler, Christian Hellriegel, Douglas S Richardson, Nicole Cayting, Nicole A. Traphagen, Wangu Mvula, Buraq Ahmed, Rinath Jeselsohn, Chongzhi Zang, Myles Brown

**Affiliations:** Department of Medical Oncology, Dana-Farber Cancer Institute, Boston, MA, 02215, USA; Department of Medicine Harvard Medical School, Boston, MA, 02215, USA; Department of Genome Sciences, University of Virginia, Charlottesville, VA, 22908, USA; UVA Comprehensive Cancer Center, Charlottesville, University of Virginia, VA, 22908, USA; Carl Zeiss Microscopy, White Plains, NY, 1060, USA; Harvard Center for Biological Imaging, Harvard University, Cambridge, MA, 02138, USA; Department of Molecular and Cellular Biology, Harvard University, Cambridge, MA, 02138, USA; Center for Functional Cancer Epigenetics, Dana-Farber Cancer Institute, Boston, MA, 02215, USA; Breast Oncology Program, Dana-Farber Brigham Cancer Center, Boston, MA, 02215, USA

## Abstract

Changes in gene expression regulated by ligand-dependent transcription factors such as estrogen receptor-α (ERα) involves the recruitment of coactivators including p300 that acetylates histone H3 at lysine 27 (H3K27ac). While H3K27ac marks active enhancers, the detailed chromatin architecture of enhancers remains unclear. Using super-resolution microscopy, we reveal distinct structural states of H3K27ac modified chromatin in response to ERα activation. In estradiol (E2)-treated cells, H3K27ac modified chromatin adopts open, elongated structures, while ERα inhibition induces compact, spherical H3K27ac modified chromatin conformations. Using MED1, a core subunit of the Mediator complex that bridges transcription factors with RNA polymerase II (Pol II), we demonstrate that larger H3K27ac structures are preferentially associated with active enhancers, whereas more compact structures show reduced MED1 association, consistent with a less active or inactive state. A constitutively active ERα mutation linked to endocrine therapy resistance in breast cancer maintains open chromatin states independent of ligand, suggesting sustained transcriptional activity. Our findings provide the first direct visualization of H3K27ac associated chromatin structural dynamics, challenging the assumption that H3K27ac modification alone is sufficient to lead to enhancer activation. By demonstrating that H3K27ac architecture is dynamically regulated by ERα, we establish a new paradigm for understanding epigenetic regulation and highlight potential therapeutic targets for endocrine therapy resistant cancers.

## Introduction

Chromatin architecture plays a fundamental role in gene regulation by dictating the accessibility of transcription machinery to DNA. Over the past decade, advances in super-resolution microscopy have revealed that chromatin is not a uniform entity but rather exists in distinct structural states that correlate with transcriptional activity [1-3]. Among the key epigenetic modifications involved in chromatin regulation, H3K27ac serves as a hallmark of active enhancers and is frequently used to identify regulatory elements associated with transcriptional activation [4]. However, while H3K27ac enrichment is widely interpreted as a proxy for enhancer activity, whether its presence alone reflects its functional state remains an open question. Growing evidence in the literature suggests that this may not always be the case. Sankar et al. (2022) used CRISPR base editing in mouse embryonic stem cells to generate pan-H3K27R mutants and directly probe the role of H3K27. They showed that while H3K27 methylation is essential for Polycomb function, H3K27 acetylation is dispensable for both gene de-repression after methylation loss and gene activation during epiblast-like differentiation. This provides proof of concept that H3K27ac does not always equate to enhancer activation or reliably indicate chromatin state and cell fate[5]

ERα is a ligand-dependent transcription factor that governs gene expression in normal estrogen responsive tissues and in approximately 70% of breast cancers [6, 7]. Upon binding to its ligand estradiol (E2), ERα recruits coactivators such as p300/CBP, which catalyzes the acetylation of H3K27 at enhancer regions to promote gene transcription [8-11]. Conversely, ERα antagonists, such as tamoxifen and fulvestrant, disrupt this process and are widely used as endocrine therapies for ER+ breast cancer [12, 13].

While ERα activity is known to regulate chromatin accessibility and enhancer function, how this regulation shapes the three-dimensional (3D) organization of H3K27ac-marked chromatin remains poorly understood. Remarkably, early ultrastructural evidence from Henri Rochefort’s 1980 electron microscopy studies revealed that physiological estradiol induces rapid chromatin decondensation in estrogen-responsive tissues, while antiestrogens like tamoxifen enforce chromatin condensation −observations that were among the first to demonstrate hormone-dependent chromatin plasticity [14]. These ligand-specific structural transitions (dispersion with ER agonists versus condensation with antagonists) suggested a link between nuclear receptor activation and global chromatin reorganization, potentially facilitating transcriptional reprogramming. However, despite decades of technological advances in chromatin visualization, these pioneering findings have seen surprisingly little follow-up investigation. Key questions remain unresolved, including whether these ultrastructural changes represent a fundamental mechanism of steroid hormone signaling, how they relate to specific transcriptional outcomes, and how they integrate with the emerging understanding of 3D genome architecture.

Recent advances in super-resolution microscopy have revolutionized our ability to study chromatin structure and transcriptional complexes, surpassing the diffraction limit of conventional light microscopy [15-18]. Traditional light microscopy (e.g., confocal), while widely used, is constrained by the diffraction properties of light, which prevent the resolution of objects closer than approximately 250 nm apart. This limitation has posed a significant challenge for investigating the intricate interactions of transcription factors (TFs) and other molecular components densely packed within the nucleus. However, the development of super-resolution microscopy, which achieves sub-50 nm resolution, has opened new avenues for exploring these complex systems [18-23]. Techniques such as Structured Illumination Microscopy (SIM) and Stochastic Optical Reconstruction Microscopy (STORM) have emerged as powerful tools, enabling researchers to visualize molecular interactions with unprecedented precision [22, 24, 25]. Despite these advances, no study has yet employed super-resolution microscopy to conduct a detailed investigation of ERα transcription complexes and its impact on chromatin architecture, leaving a critical gap in our understanding of its mechanistic role in gene regulation.

Notably, Boettiger et al. (2016) employed STORM to show that chromatin is organized into discrete nanoscale domains, which can be categorized as active, repressed, or inactive states based on their structural properties [15]. Their study demonstrated that active chromatin regions exhibit larger, more dispersed structures, whereas repressed chromatin is characterized by compact, dense formations. These findings align with other work showing that chromatin organization is dynamically regulated in response to transcriptional activation [16, 17], and that epigenetic modifications such as H3K27ac play a critical role in shaping chromatin topology [19, 20, 22].

Unlike previous studies that rely on FISH-based assays −which require DNA denaturation and disrupt native nuclear architecture− our work leverages super-resolution microscopy to directly visualize chromatin structure in fixed cells. This approach preserves the physiological context of the nucleus, enabling the study of chromatin dynamics under conditions that closely mimic the cellular environment. Furthermore, our study uniquely investigates the regulation of H3K27ac modified chromatin in response to the activation of a transcription factor, ERα, via its natural ligand E2. This focus on a physiologically relevant stimulus provides novel insights into the dynamic interplay between transcriptional activation and chromatin structure [21, 24, 26, 27].

Given the critical role of chromatin structure in enhancer function, in this study we employ super-resolution SIM and STORM to visualize the nanoscale organization of H3K27ac modified chromatin in response to ERα activation and inhibition, revealing that ERα-induced chromatin changes coincide with distinct structural alterations. By systematically analyzing H3K27ac under various conditions, including E2 stimulation, ER antagonists, and clinically relevant ERα mutations linked to endocrine therapy resistance, we establish a direct connection between ERα activity and the spatial organization of H3K27ac-marked chromatin. To further assess enhancer activation, we quantified the association of H3K27ac structures with MED1, a core Mediator subunit that bridges transcription factors to RNA polymerase II and marks active enhancers. Our results show that larger and non-spherical H3K27ac structures associate significantly more with MED1, thereby challenging the conventional view that H3K27ac enrichment alone defines enhancer activation. Instead, our findings highlight chromatin topology as a key regulator of transcription, advancing our understanding of ERα-mediated epigenetic regulation and introducing a new paradigm for studying enhancer activity through chromatin structure.

## Results

### ER Activation Enhances Proximity to H3K27ac-Modified Chromatin

Previous genome-wide studies using ChIP-seq have demonstrated that ERα activation leads to increased level of H3K27ac, mediated by the recruitment of histone acetyltransferases (HATs) such as p300 to specific gene promoters and enhancers. This epigenetic modification is associated with an open chromatin state, facilitating the binding of transcription factors and the transcription machinery [28-30]. However, these bulk genomic assays lack the spatial resolution to capture the dynamic structural changes of chromatin at the single-cell level.

To address this limitation, we employed super-resolution SIM to visualize the spatial organization of ERα and H3K27ac in fixed MCF7 cells. While confocal microscopy provides a global nuclear signal, it lacked the resolution to distinguish specific spatial relationships between ERα and H3K27ac within the densely packed nuclear environment (Fig. 1A). In contrast, SIM enabled us to resolve distinct structural changes in H3K27ac domains with significantly higher precision (Fig. 1B). Using a data-driven thresholding approach to analyze the SIM data [31], we quantified the correlation between ERα and H3K27ac signal intensity across a large dataset, comprising 300–700 cells and more than 100,000 H3K27ac domains (Fig. 1C, D, Fig. S1A-C, Table S1).

**Figure 1.**
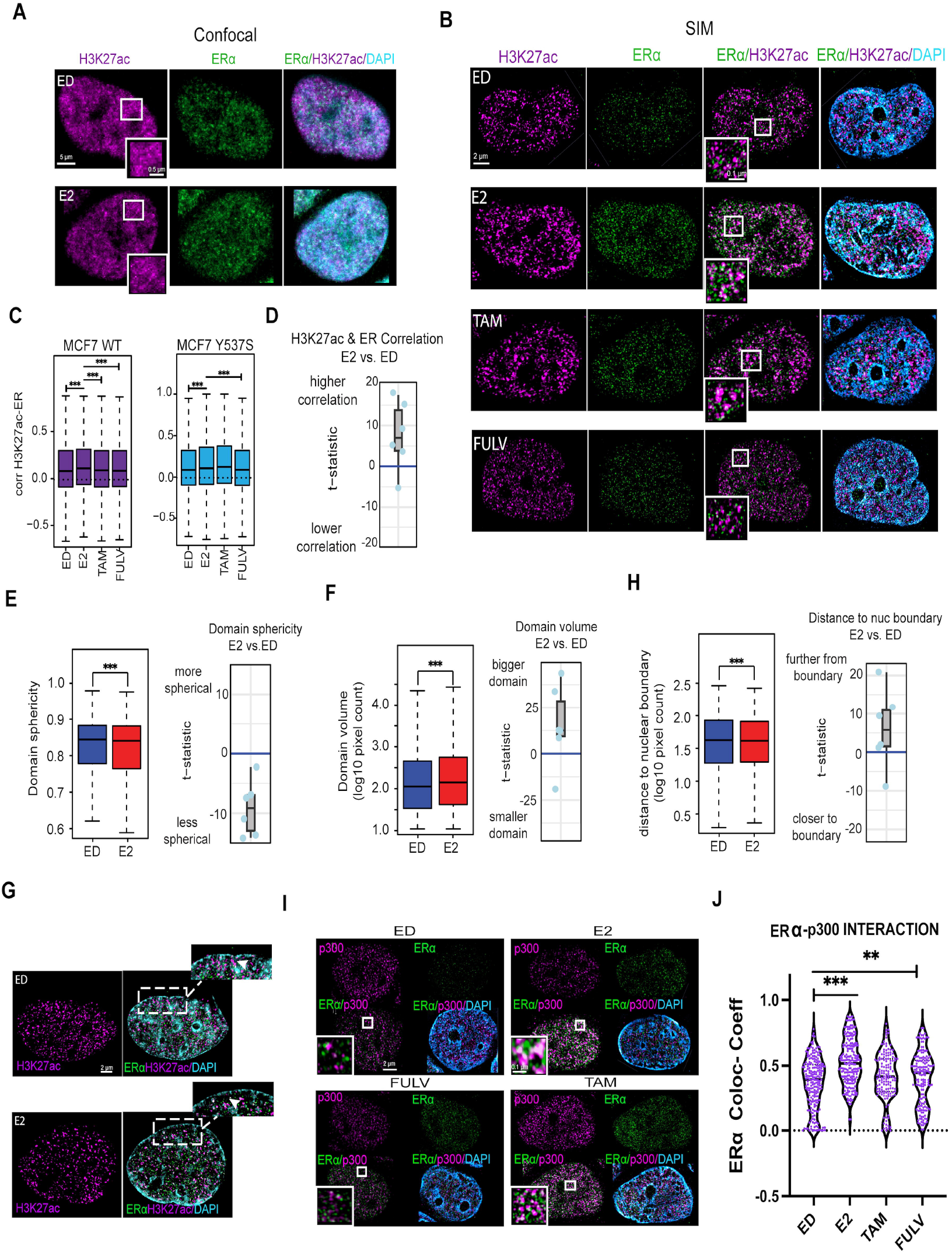
ER Activation Enhances Proximity to H3K27ac-Modified Chromatin. **(A)** Confocal images of MCF7 cells probed for H3K27ac (magenta) and ERα (green) under estrogen-deprived (ED) and E2-treated (10 nM) conditions, highlighting the resolution limit of conventional microscopy. **(B)** Super-resolution 3D-SIM images of MCF7 cells under ED, E2 (10 nM), Tam (100 nM), and Fulv (100 nM) conditions, demonstrating enhanced proximity between ERα and H3K27ac in E2-treated cells. **(C)** Boxplots of pixel-level correlation coefficients between ERα and H3K27ac signal intensities in H3K27ac domains for WT and Y537S mutant MCF7 cells under ED, E2, Tam, and Fulv conditions. Statistics (one-side student T-test) were provided for E2 compared with other conditions: *P < 0.01, **P < 0.001, ***P < 0.0001. Quantification was performed on 300-700 cells per treatment group. **(D)** Boxplot with overlaid data points showing the t-statistic from each data set. The t-statistic quantifies the difference in correlation between H3K27ac and ER signal intensity in E2 compared to ED. Positive and negative scores indicate higher and lower correlation, respectively. **(E)** Domain sphericity of the H3K27ac domains in ED vs. E2 (blue and red, respectively) in WT MCF7 cells. Boxplot of t-statistics shows sphericity differences (E2 vs. ED); negative scores indicate reduced sphericity in E2-treated cells. **(F)** Boxplots showing the volume of H3K27ac domains in under ED and E2-treated conditions, with t-test showing higher domain volume in E2. Boxplot of t-statistics shows volume differences (E2 vs. ED); positive scores indicate larger volumes in E2. **(G)** 3D-SIM images of MCF7 cells under ED, E2. Zoom-in panels highlight differences in H3K27ac positioning. In ED, arrows indicate H3K27ac foci localized closer to the nuclear periphery, whereas in E2, H3K27ac appears redistributed further toward more internal nuclear regions. **(H)** Boxplots showing the distance from H3K27ac domains to nearest nuclear boundary in ED and E2-treated conditions, with t-test showing the greater distance in E2 from the nuclear boundary. Boxplot of t-statistic scores for the difference of H3K27ac domain to the nearest nuclear boundary (E2 vs. ED). Positive scores indicate in E2, H3K27ac domains are farther from the boundary. All quantifications were performed in 300-700 cells per treatment group. *P < 0.01, **P < 0.001, ***P < 0.0001. **(I)** 3D-SIM images of MCF7 cells probed for p300 (magenta) and ERα (green) under ED, E2, Tam, and Fulv conditions. **(J)** Violin plots of colocalization coefficients between p300 and Erα under different conditions. Quantification was performed in n >140 cells per treatment group. *P < 0.01, **P < 0.001, ***P < 0.0001.

In estrogen-deprived (ED) cells, ERα showed moderate correlation with H3K27ac. However, upon E2 treatment, this correlation increased significantly (P < 0.001), indicating ERα-driven chromatin activation. Conversely, treatment with the ERα antagonists fulvestrant (Fulv) and tamoxifen (Tam), reduced this association, consistent with their role in inhibiting ERα activity (Fig. 1C, and Fig. S1D, Table S1). Intriguingly, using a doxycycline-inducible MCF7 cell line expressing HA-tagged Y537S ERα, a clinically relevant mutation known for its ligand-independent activity, we observed that Y537S ERα maintained a correlation with H3K27ac regardless of E2 or antagonist treatment. However, this correlation was weaker compared to wild-type (WT) ERα (Fig. 1C and Fig. S1E, F, Table S1). This finding suggests that while the ERα Y537S mutation is sufficient to drive hormone independent breast cancer growth, it may have properties that are distinct from WT ERα activated by E2. This is consistent with prior work demonstrating allele-specific transcriptional programs regulated by ERα [32].

To further investigate the structural changes in H3K27ac domains, we analyzed their morphology under different ERα activation states. We first observed that, in E2-treated cells, H3K27ac domains exhibited significantly lower sphericity compared to ED condition, suggesting a more elongated and irregular shape. In contrast, H3K27ac domains tend to be more spherical in ED cells (Fig. 1E, S2A, Table S1). Additionally, quantitative analysis (300–700 cells per analysis, Table S1), revealed that the H3K27ac domains was significantly larger in E2-treated cells compared to ED, Tam, and Fulv conditions (Fig. 1F, S2B). These findings suggest that the observed structural changes may be linked to the activation state of H3K27ac, as previous studies using STORM have demonstrated that active chromatin regions tend to adopt less condensed, more dispersed structures. In contrast, inactive or repressed chromatin is typically characterized by compact, densely packed formations [15].

Furthermore, majority of H3K27ac domains in E2-treated cells were positioned farther from the nuclear boundary, suggesting a shift from peripheral lamina-associated domains to central, transcriptionally active regions (Fig. 1G, H, S2C, Table S1) [33].This repositioning is consistent with the role of ERα in promoting chromatin accessibility and enhancer activation and H3K27ac engagement in active transcription, as central nuclear regions are enriched in transcriptionally active chromatin [33, 34].

Next, we investigated the interaction between ERα and p300 in response to the various ER ligands (Fig. 1I). E2 treatment enhanced the association of ERα with p300, while Fulv and Tam treatments diminished this interaction (Fig. 1I, J, Table S2). In Y537S mutant expressing cells, p300 association with ERα persisted regardless of ligand availability, though at lower levels compared to E2-treated WT cells. Notably, Fulv, but not Tam, reduced this association in Y537S cells (Fig. S2 D, E, Table S2). These results are consistent with the observed ERα-H3K27ac correlation and highlight the role of p300 in mediating ERα-dependent chromatin modification [32, 35, 36].

### H3K27ac Structural Changes Correlate with ERα Activation

Given that previous studies have established a link between epigenetic modifications and chromatin architecture activation state [15], we employed higher-resolution STORM microscopy to further resolve structural changes in H3K27ac in response to ERα activation and chromatin remodeling. While SIM imaging provides substantially greater resolution than conventional confocal microscopy (∼100 nm lateral, ∼500 nm axial), it remains limited in resolving finer chromatin features. By contrast, STORM offers markedly improved spatial resolution (∼30 nm XY, ∼60 nm Z), enabling more accurate delineation of chromatin domain structures and more precise identification of distinct chromatin morphologies, thereby strengthening our structural inferences.

Initially, we applied two-color 2D STORM microscopy. In E2-treated cells, H3K27ac structures appeared elongated and less condensed, while ED cells exhibited punctate, compact formations (Fig. S3A). Although 2D STORM provided valuable insights into H3K27ac domain morphology, it lacked the depth resolution required to fully capture the 3D structural dynamics of chromatin. Additionally, 2D imaging is prone to optical illusions caused by overlapping structures and projection artifacts, which can obscure the true spatial organization of chromatin.

To overcome the limitations of 2D imaging, we transitioned to 2-color 3D STORM microscopy, which provided a more detailed and nuanced picture of chromatin organization in intact cells. Using 3D STORM, we observed that E2 deprivation induced globular H3K27ac structures, whereas E2 stimulation resulted in elongated, dispersed conformations (Fig. 2A, n> 25 cells; Table S3). To confirm the ERα activation state upon E2 treatment, we probed for both ERα and p300 and imaged the cells using 3D STORM. As expected, ERα correlated strongly with p300 in E2-treated cells (Fig. S3B), confirming that ERα is activated in E2-treated cells.

**Figure 2.**
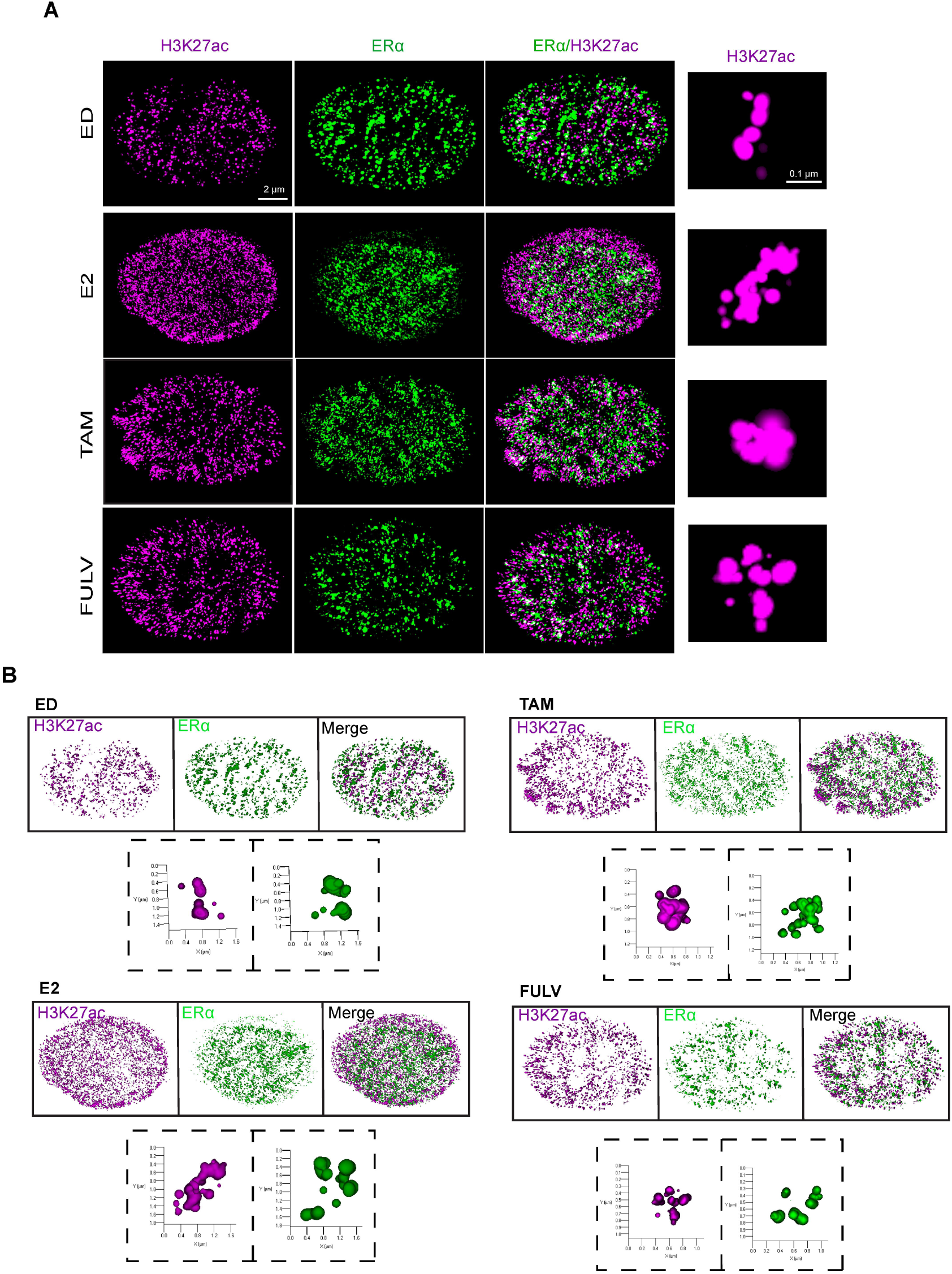
H3K27ac Structural Changes Correlate with ERα Activation. **(A)** 2-color 3D STORM images of MCF7 cells treated with E2 (10 nM), Tam (100 nM), and Fulv (100 nM). Tam and Fulv treatments show more compact H3K27ac structures compared to the elongated structures in E2-treated cells. **(B)** 3D renderings of STORM images from Fig. 2A, indicating elongated and compact structures for each condition.

Next, we treated cells with two ERα antagonists commonly used in endocrine therapy, Tamoxifen and Fulvestrant, and performed 3D-STORM to examine H3K27ac structural changes induced by these treatments. Similar to E2-deprived cells, and in contrast to E2-treated cells, most H3K27ac domains appeared more spherical and compact rather than elongated and dispersed (Fig. 2A, n ≥15 cells; Table S3). Importantly, these phenotypes—although broadly categorized as elongated/irregular versus spherical—should not be viewed as strictly binary states. Instead, they likely represent condition-dependent remodeling of enhancer architecture along a continuum, rather than discrete markers of activity.

To further characterize and quantify the structural states of H3K27ac, we first employed AI-based rendering tools to classify individual chromatin domains into distinct morphological categories. In E2-treated cells, H3K27ac predominantly formed “open-like” structures, characterized by larger volumes and irregular shapes (Fig. 2B, Movie 1, 2). In contrast, ED cells and those treated with ERα antagonists (Tam and Fulv) exhibited “closed-like” structures, with smaller volumes and higher sphericity (Fig. 2B, Movie 3-8). Together, these imaging results suggest that ERα activity influences H3K27ac chromatin architecture, producing distinct structural conformations under different hormonal and therapeutic conditions. While these observations point to a potential relationship between domain morphology and enhancer activity, systematic quantification is required to define the extent and significance of these structural changes.

### Structural Changes in H3K27ac Are Dependent on ERα Coactivators

Next, to determine the role of ERα coactivators in regulating H3K27ac structural dynamics, we systematically inhibited key components of the ERα transcriptional machinery. Knockdown of NCOA3, a critical ERα coactivator that links ERα to p300 [37], abolished the formation of elongated H3K27ac structures in E2-treated cells, resulting in chromatin domains that resembled those observed in ED or antagonist-treated conditions (Fig. 3A, S3C-E, n>20 cells; Table S3). Similarly, pharmacological inhibition of p300 using A-485 prevented the transition to dispersed chromatin states (Fig. 3B, C, S3F), further underscoring the essential role of p300-mediated acetylation in chromatin remodeling. Notably, when we measured global H3K27ac intensity, we found no significant differences across the various experimental conditions, except in E2-treated cells, where H3K27ac intensity was higher on average (Fig. S4A, Table S3). This suggests that while ERα activation enhances H3K27ac intensity, the baseline levels of H3K27ac in majority of states remain relatively stable. Furthermore, the structural changes observed appear to be influenced by the activation state of ERα, rather than by global shifts in H3K27ac levels. This highlights a potential role for ERα in modulating chromatin architecture through localized, rather than global, changes in H3K27ac.

**Figure 3.**
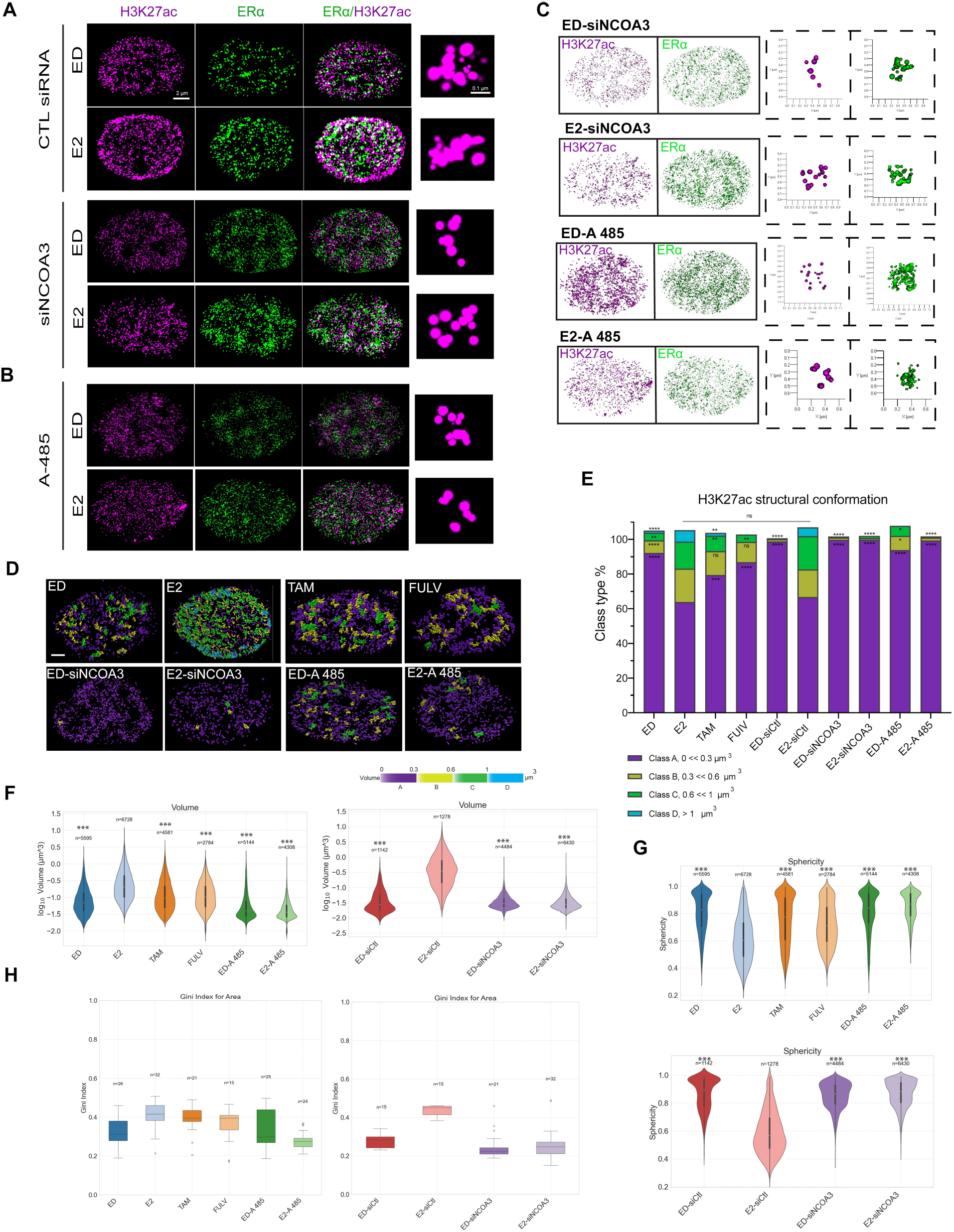
Structural Changes in H3K27ac Are Dependent on ERα Coactivators. **(A, B)** 3D STORM images of MCF7 cells under (A) control siRNA or NCOA3 siRNA and (B) p300 inhibitor A485 (80 nM) conditions. The elongated H3K27ac structures observed in E2-treated control siRNA cells were lost upon NCOA3 knockdown or p300 inhibition. **(C)** Renderings of STORM images from (A, B), highlighting elongated versus compact H3K27ac structures in cells lacking NCOA3 or p300 activity. **(D)** Structural maps generated from rendered images in Figures 2 and 3 (A, B), classifying H3K27ac-modified chromatin into four volume classes (purple, yellow, green, blue; low to high volume). **(E)** Distribution of volume classes across conditions. E2-treated cells displayed more high-volume elongated structures, whereas Tam, Fulv, NCOA3 siRNA, and A485 treatments shifted the distribution toward low-volume compact structures. *P < 0.01, **P < 0.001, ***P < 0.0001. **(F, G)** Violin plots of H3K27ac domain volumes (F) and sphericity (G) across conditions. Kruskal–Wallis ANOVA followed by Mann–Whitney U test with Bonferroni correction. *P < 0.01, **P < 0.001, ***P < 0.0001. **(H)** Gini index of H3K27ac volume heterogeneity across conditions. Kruskal–Wallis test followed by Mann–Whitney U test with Bonferroni correction. All quantifications were performed in 15–35 cells, with >1200 H3K27ac domains analyzed per treatment group.

The dependence of H3K27ac structural changes on NCOA3 and p300 highlights the critical role of ERα coactivators in shaping chromatin architecture. In the absence of functional NCOA3 or p300, ERα is unable to induce the structural changes observed, leading to a shift toward more compact chromatin states.

This is consistent with previous studies showing that coactivators including NCOA3 and p300 facilitate the recruitment of transcriptional machinery to ERα target gene expression [7, 34]. Importantly, the loss of elongated H3K27ac structures upon coactivator inhibition demonstrates that these coactivators are essential for ERα-mediated chromatin remodeling. Together, these findings confirm that the structural changes in H3K27ac are driven by ERα transcriptional activity and its associated coactivators. The observed dependence on NCOA3 and p300 provides a direct link between ERα signaling, chromatin architecture, and epigenetic modifications.

To quantify these structural changes observed in figures 2 and 3, we classified H3K27ac domains into four volume-based categories and generated distribution maps for each condition (Fig. 3D). E2-treated cells showed a higher percentage of larger volume distributions (yellow, green, and blue) compared to ED and antagonist treatments (purple). Additionally, silencing NCOA3 or inhibiting p300 shifted the percentage of volume distribution toward lower volumes (Fig. 3E, Fig. S4B-E, Table S3, Movie 9-12). These results further highlight the critical role of ERα-coactivator complexes in regulating H3K27ac-mediated structural dynamics. We then quantified all H3K27ac domain volumes across different conditions and along all three X, Y, and Z axes. Consistently, the volume in the active ERα condition (E2 treatment) was higher and less spherical compared to ED, antagonist, and inactive coactivator conditions (Fig. 34F-G, S4F-I).

To assess the heterogeneity of H3K27ac structures across different conditions, we performed a Gini index analysis, a statistical measure of inequality often used to evaluate structural diversity. E2-treated cells exhibited the highest Gini index, indicating greater heterogeneity in chromatin domain sizes and shapes, consistent with dynamic chromatin remodeling during transcriptional activation (Fig. 3H). In contrast, ED cells and those treated with ERα antagonists showed lower Gini indices, reflecting more uniform chromatin organization associated with transcriptional repression. This suggests that the structural heterogeneity observed in E2-treated cells is driven by ERα activity and its ability to recruit coactivators, which promote diverse chromatin conformations. Conversely, inhibition of p300 with A-485 resulted in the lowest Gini index, indicating that p300 activity is essential for generating the structural diversity associated with active chromatin (Fig. 3H) and as a critical regulator of chromatin dynamics in cancer [34].

These findings highlight the dynamic nature of chromatin architecture in response to transcriptional stimuli and establish a quantitative framework for understanding how epigenetic modifications, such as H3K27ac, contribute to gene regulation. The observed heterogeneity in chromatin structures highlights the complexity of enhancer regulation and suggests that chromatin topology plays a critical role in determining transcriptional outcomes. Importantly, the expression of specific epigenetic markers alone is not sufficient to fully explain these regulatory mechanisms [38-40]. This notion is further supported by studies that have delineated the organizational principles of 3D genome architecture, identifying topological domains in mammalian genomes as key regulators of chromatin interactions [17, 27, 41, 42]. Together, these insights emphasize the importance of integrating both epigenetic modifications and spatial chromatin organization to fully understand gene regulation.

### H3K27ac Structural Changes Reflect Chromatin Accessibility

Given that our observations align with previous STORM studies showing that active chromatin domains are typically larger and more dispersed, whereas inactive domains are compact and more spherical [15], we next asked whether H3K27ac structures are linked to enhancer activation state. To address this, we probed H3K27ac domains with MED1, a known ER coactivator and mediator of RNA Polymerase II. MED1 enrichment is widely used in techniques such as ChIP-seq as a marker of active enhancer states [43-45]. We therefore examined MED1 localization in both E2-treated and E2-deprived cells and analyzed its correlation with H3K27ac morphology.Larger, elongated H3K27ac structures were strongly enriched for MED1, consistent with active enhancer elements, whereas smaller, spherical domains showed significantly less MED1 association, consistent with inactive enhancers (Fig. 4A,B, n>65 cells; Table S4). These findings are in line with previous reports that larger chromatin volumes represent accessible and transcriptionally active regions, whereas smaller, compact structures reflect inaccessible states [15]. Importantly, the overall correlation of MED1 with both H3K27ac and ER was significantly higher in E2-treated than in E2-deprived cells, demonstrating that enhancer activation and H3K27ac reorganization are orchestrated by ERα signaling (Fig. S5A–C; Table S4).

**Figure 4.**
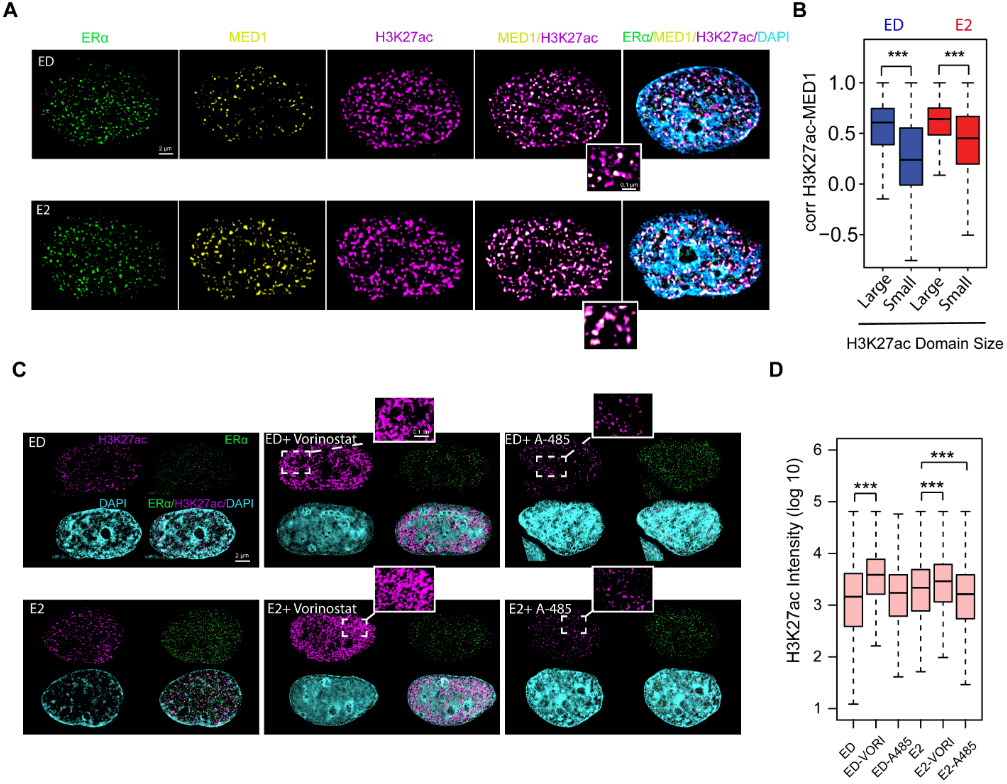
H3K27ac Structural Changes Reflect Chromatin Accessibility. **(A)** Super-resolution 3D-SIM images of MCF7 cells cultured under ED or E2 (10 nM) conditions, stained for H3K27ac, ER, and MED1. **(B)** Boxplots of pixel-level correlation coefficients between MED1 and large versus small H3K27ac structures. Larger H3K27ac domains show significantly stronger association with MED1, consistent with an active enhancer state. ***P < 0.0001. Quantification was performed on 65–150 cells per treatment group. **(C)** Cells treated with the histone deacetylase (HDAC) inhibitor vorinostat or the histone acetyltransferase (HAT) inhibitor A-485. Vorinostat treatment induced a robust global increase in H3K27ac signal intensity, whereas A-485 treatment shifted the distribution toward puncta-like foci. **(D)** Quantification of H3K27ac signal intensity under the indicated conditions. Vorinostat treatment produced a marked global increase in H3K27ac, although high signal density precluded reliable segmentation of individual domains for structural analysis. A-485 treatment revealed a shift in H3K27ac structures between active and inactive acetylation states. Quantification was performed on 50–260 cells per treatment group.

As a complementary approach, we perturbed global histone acetylation to further probe the link between H3K27ac structure and activity. Treatment with the histone deacetylase (HDAC) inhibitor vorinostat resulted in a robust global increase in H3K27ac signal intensity, whereas treatment with the histone acetyltransferase inhibitor A-485 produced a shift toward numerous puncta-like foci (Fig. 4C). Quantification confirmed the global increase in H3K27ac after vorinostat, though the high density of signals precluded confident segmentation of individual domains for structural analysis (Fig. 4D, n>50 cells; Table S5). These pharmacological manipulations reinforce that H3K27ac morphology is sensitive to the acetylation state: hyperacetylation drives widespread signal with diffuse morphology, while inhibition of acetylation yields fragmented punctate structures.

Together, these findings demonstrate that transitions between elongated and compact H3K27ac domains reflect enhancer activity state. MED1 association and pharmacological perturbation both support the conclusion that “open,” accessible H3K27ac-modified chromatin is structurally and functionally distinct from its “closed,” inaccessible counterpart. These H3K27ac states are governed by ERα and its coactivators, establishing a mechanistic link between ERα signaling, enhancer activation, and chromatin architecture.

### Ligand-Independent ERα Mutations Maintain Active Chromatin Structures

To investigate the chromatin regulatory mechanisms driving endocrine resistance, we focused on the Y537S ERα mutant [32, 35, 36, 46]. We employed 3D STORM imaging to analyze H3K27ac structures in the presence and absence of E2 in cells expressing the Y537S mutant. Strikingly, even under ED conditions, cells expressing Y537S exhibited open, elongated H3K27ac structures (Fig 5. A, B, n>15 cells; Table S3). Although these structures resembled those observed in E2-treated WT cells, key differences were apparent. First, the overall frequency of elongated structures in Y537S cells was lower than in WT-E2 cells but higher than in WT-ED cells (Fig. 5A-D). Second, the elongated H3K27ac structures in Y537S cells were shorter in length, indicating that the structural remodeling induced by Y537S differs from that induced by E2 in WT cells (Fig. S6A). These structural differences were further reflected in distinct characteristics such as sphericity, volume, area, and Gini index quantifications observed between Y537S and E2-treated WT cells (Fig 5E-F, Fig. S6B-C, Table S3).

**Figure 5.**
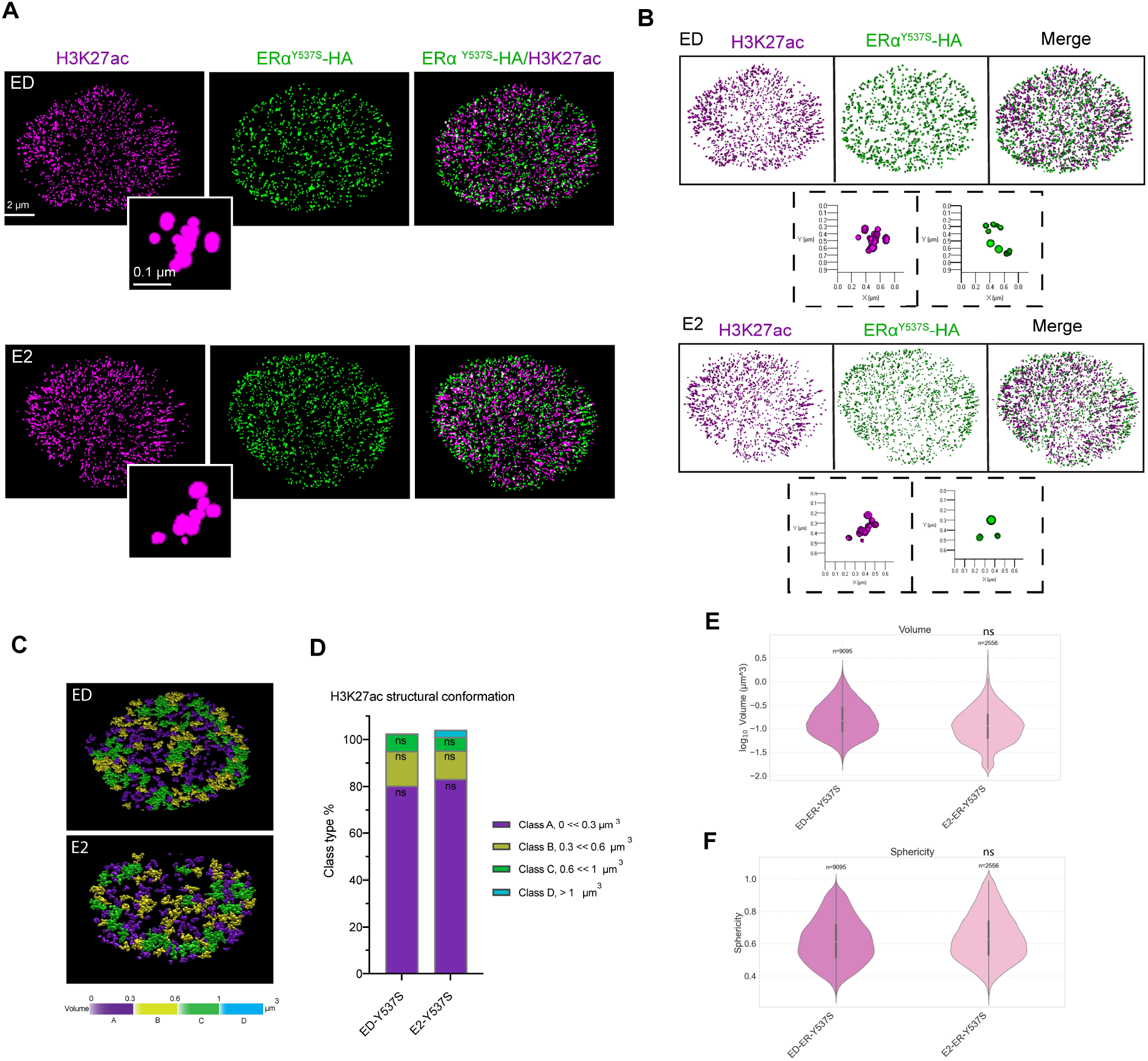
Ligand-Independent H3k27ac Chromatin Modification in Y537S ERα Mutant Cells. **(A)** 3D-STORM images of MCF7 Y537S cells under ED and E2-treated conditions, showing “open” structures independent of ligand. **(B, C)** 3D renderings (B) and structural maps (C) of chromatin volume classes (yellow/green/blue = high; purple = low), showing “closed” vs. “open” structures. **(D)** Distribution percentage of each volume class for ED and E2 conditions in Y537S cells, showing no significant change. **(E, F)** Violin plots of H3K27ac domain volumes (E), and sphericity (F), for ED and E2 conditions in Y537S cells. Kruskal-Wallis ANOVA shows no significant difference between ED and E2 conditions.

This is consistent with previous findings showing that Y537S regulates allele-specific transcriptional programs [32]. Together, these observations suggest that the Y537S mutation promotes chromatin accessibility in a ligand-independent manner, consistent with its constitutive transcriptional activity. These results offer mechanistic insights into how Y537S may drive endocrine resistance by maintaining an active chromatin state independent of hormonal signaling.

Quantitative analysis revealed that the volume and sphericity of H3K27ac domains in Y537S mutant cells were comparable to those in E2-treated WT cells, further supporting the idea that this mutation sustains an active chromatin state (Fig. 5C-F, S6A-B, Table S3). Notably, there was no significant change in global H3K27ac signal intensity in the presence or absence of ligand (Fig. S6C), suggesting that the observed structural changes in chromatin remodeling and H3K27ac organization are driven independent of ligand. Additionally, the Gini index, a metric for structural heterogeneity, revealed that Y537S mutant cells displayed moderate heterogeneity, in contrast to the uniform chromatin organization typically seen in ED or antagonist-treated WT cells (Fig. S6D).

The ability of the Y537S mutation to maintain active chromatin structures in the absence of ligand highlights its role in driving endocrine resistance. This is consistent with previous studies showing that ligand-independent ERα mutations can maintain transcriptional activity even in the absence of estrogen [46, 47]. Our findings extend this model by showing that these mutations also promote an open chromatin state, facilitating continuous transcriptional activity even under hormone-deprived conditions. These results provide mechanistic insights into how ERα mutations contribute to endocrine resistance by maintaining an active chromatin state. The observed structural changes in H3K27ac domains suggest that these mutations alter the epigenetic landscape to active chromatin, enabling persistent gene expression and tumor progression. This underscores the potential of targeting chromatin modification and remodeling pathways in overcoming resistance to endocrine therapies.

## Discussion

Our study leverages the super resolution of STORM microscopy to provide the first direct visualization of H3K27ac modified chromatin structural dynamics in response to ERα activity. By revealing that H3K27ac adopts distinct structures that reflect enhancer activity, we refine the conventional view that H3K27ac enrichment alone defines enhancer activity. We demonstrate that chromatin architecture plays a critical role in determining the functional state of enhancers, offering a new paradigm for understanding transcriptional regulation.

Our study bridges a critical gap in epigenetic research by revealing that H3K27ac-marked chromatin exists in structurally and functionally distinct open/closed states. These findings provide mechanistic clarity to early observations by Rochefort’s group (1980), who first documented estrogen-induced chromatin decondensation using electron microscopy but lacked the tools to link ultrastructural changes to specific epigenetic marks or transcriptional outcomes [14]. Where their work identified ligand-dependent global chromatin reorganization (estradiol-induced dispersion vs. tamoxifen-driven condensation), we now demonstrate that these dynamics are encoded at the nanoscale through H3K27ac architectural switching.

In this work, we used two-color 3D-STORM microscopy to resolve nanoscale chromatin structures in intact cells. Unlike bulk assays such as ChIP-seq, which provide population-averaged data, SIM and STORM enable single-molecule localization with nanometer precision, allowing us to directly observe the spatial organization of H3K27ac in response to ERα activation [21]. This approach revealed that ERα-driven chromatin remodeling is accompanied by dramatic changes in H3K27ac domain morphology, including increased volume, reduced sphericity, and repositioning away from the nuclear periphery. These findings provide unprecedented insights into the dynamic nature of chromatin architecture and its role in gene regulation.

Our discovery that H3K27ac modified chromatin can fold into both open and closed states has broad implications for understanding enhancer function. While open H3K27ac structures correlate with active transcription, closed structures are associated with transcriptional repression, suggesting that chromatin topology is a key determinant of enhancer activity. Using MED1 as a functional marker, we find that elongated, dispersed H3K27ac domains are preferentially enriched for active enhancer components, whereas compact, spherical domains show little MED1 association and reflect inactive states. This is consistent with prior work linking larger, open chromatin structures to accessibility and smaller, constrained conformations to inaccessibility [15]. H3K27ac expression alone does not always denote enhancer activation or reliably reflect chromatin state, as highlighted by prior studies [5]. Notably, functional activity was revealed only when H3K27ac morphology was considered in the context of MED1 and ERα association, which was markedly higher under E2 stimulation. This underscores ERα’s central role in enhancer activation and chromatin remodeling. Pharmacological perturbation further supports this model. HDAC inhibition with vorinostat drove a diffuse global increase in H3K27ac, while acetyltransferase blockade with A-485 produced fragmented puncta-like structures. These contrasting morphologies demonstrate that the balance of acetylation not only dictates overall H3K27ac levels but also shapes its spatial organization, linking domain architecture directly to enhancer activity.

The use of STORM microscopy also elucidated the role of ERα coactivators in regulating H3K27ac structural dynamics. We demonstrated that inhibition of p300 or knockdown of NCOA3 prevents the formation of open chromatin states, underscoring the importance of coactivator activity in chromatin remodeling. These findings align with previous literature showing that p300-mediated histone acetylation regulates ERα activity, highlighting the role of p300-dependent chromatin remodeling in transcriptional activity and cell fate decisions [48]. These insights not only advance our understanding of ERα-mediated gene regulation but also suggest that coactivators could be viable therapeutic targets for disrupting chromatin architecture in endocrine-resistant cancers.

Furthermore, the phase separation condensate framework has profoundly influenced our understanding of nuclear organization, offering compelling explanations for chromatin compartmentalization and transcription factor dynamics [49]. However, the field must critically examine whether current methodologies adequately capture the complexity of these processes in living cells. While studies of estrogen receptor (ER) signaling [50-52] and epigenetic regulation (e.g., H3K27ac-mediated enhancer hubs or HP1-driven heterochromatin) present convincing *in vitro* evidence for phase separation, their *in vivo* conclusions often rely disproportionately on puncta observed via conventional microscopy [53-55]. This methodological gap creates a concerning disconnect: although purified components may exhibit liquid-like properties in reductionist systems, their behavior in the crowded, structured nuclear environment likely differs substantially. These concerns are compounded by the field’s heavy reliance on puncta as primary evidence for *in vivo* phase separation. Super-resolution techniques, including our findings, reveal that presumed “condensates” exhibit more defined architectures or stable interactions than predicted by liquid-phase separation (LLPS) models [3, 15]. Critically, diffraction-limited microscopy cannot distinguish between true liquid condensates, stable protein-DNA complexes, or imaging artifacts -all of which may appear as similar punctate structures. Moving forward, the field must integrate high-resolution imaging to refine our understanding of nuclear organization, ensuring that the condensate hypothesis is applied judiciously rather than as a default explanation for all punctate structures. This does not dismiss the importance of phase separation in gene regulation but calls for more rigorous validation. Techniques such as STORM, PALM, and electron microscopy are essential to distinguish true condensates from structured assemblies and to bridge the gap between *in vitro* models and *in vivo* complexity.

Finally, our study underscores the clinical importance of chromatin architecture in breast cancer. We show that the ERα Y537S mutation preserves open chromatin structures resembling those induced by E2-bound WT ERα, yet with distinct features that may explain its allele-specific activity [32, 35]. These findings reveal how chromatin architecture contributes to the establishment of epigenetic states in endocrine-resistant cells and offer a mechanistic basis for therapy resistance. Persistent open chromatin driven by ERα mutants enables sustained transcriptional activity despite therapeutic pressure, consistent with prior studies linking activating ESR1 mutations to hormone-resistant metastatic breast cancer and allele-specific chromatin recruitment to resistance mechanisms [32, 46, 47]. By directly connecting chromatin architecture to transcriptional output, our work highlights chromatin remodeling pathways—not merely histone marks or protein levels—as actionable vulnerabilities, opening new directions for therapeutic intervention in ERα mutant breast cancer.

In conclusion, this work establishes a new framework for studying enhancer activity through the lens of chromatin architecture. By combining super-resolution microscopy with functional assays, we have uncovered the dynamic structural states of H3K27ac modified chromatin and their role in transcriptional regulation. These findings offer mechanistic clarity to early observations by Vic et al. (1980), that first described estrogen-induced chromatin decondensation [14]. These insights not only enhance our understanding of epigenetic regulation but also open new avenues for therapeutic intervention in hormone-dependent cancers. Future studies could explore the broader applicability of these findings to other transcription factors and cancer types, as well as the potential for targeting chromatin architecture in precision medicine.

## Data availability

Raw image stacks, after reconstruction but prior to further processing, are included with the manuscript (main figures and Supplementary Information). Due to file size (>12 TB), the full unprocessed raw datasets are available from the corresponding author upon reasonable request.

## Code availability

All image analysis pipelines and custom scripts used in this study are publicly available on GitHub at https://github.com/zang-lab/3DSIManalysis.

## Acknowledgements

We thank the Harvard Center for Biological Imaging (RRID:SCR_018673) for their infrastructure and expert assistance. We also thank Molecular Imaging Core (MIC) at Dana-Farber Cancer Institute for their assistance. This work was supported in part by grants to MB from the Ludwig Center at Harvard Medical School, the Breast Cancer Research Foundation, and Susan G. Komen. CZ is partially supported by NIH grant R35GM133712 and a Pinn Scholar Award from University of Virginia School of Medicine.

