## Supplementary figures and images for "Super-resolution microscopy reveals distinct epigenetic states regulated by estrogen receptor activity"

### Supplementary Figure 1

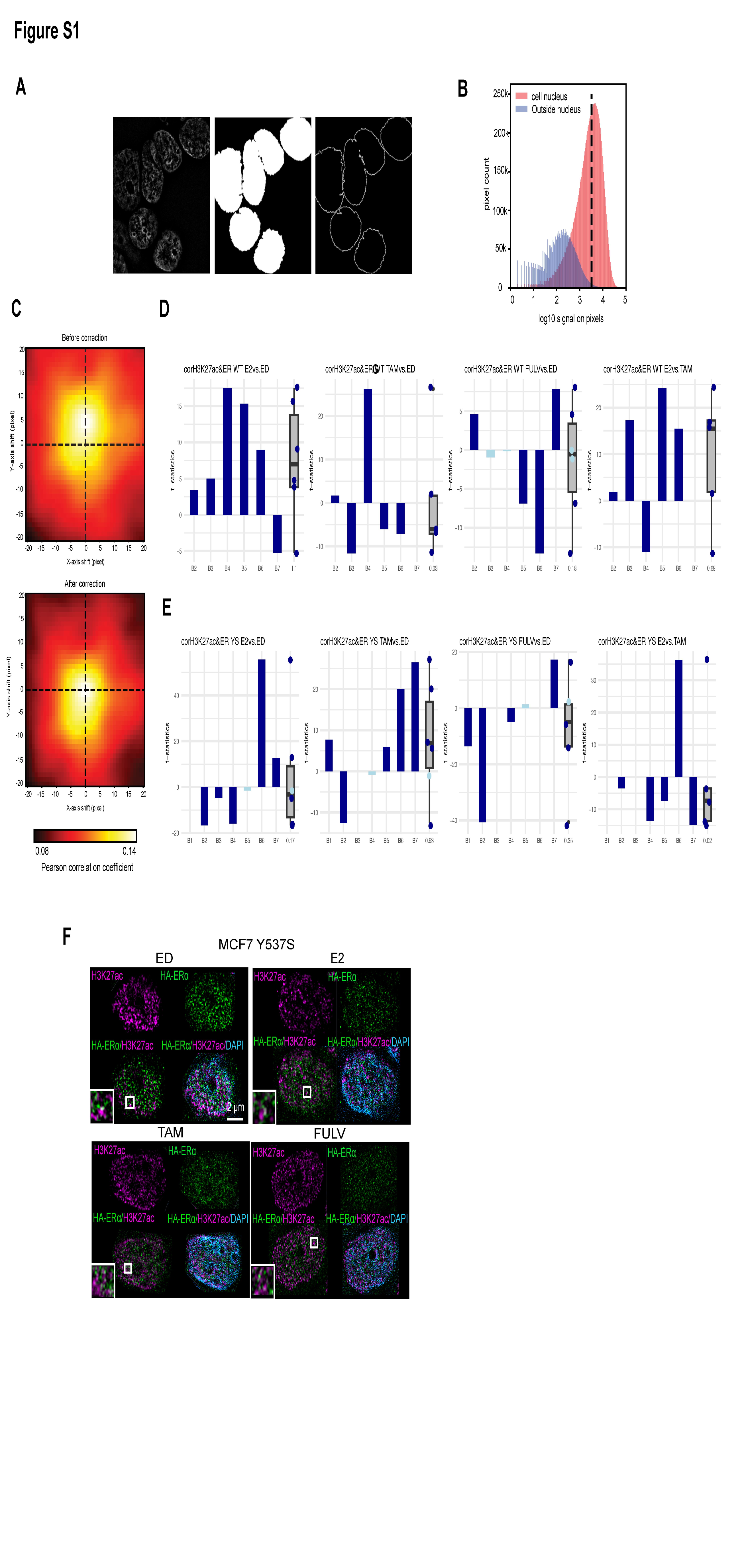

### Supplementary Figure 2

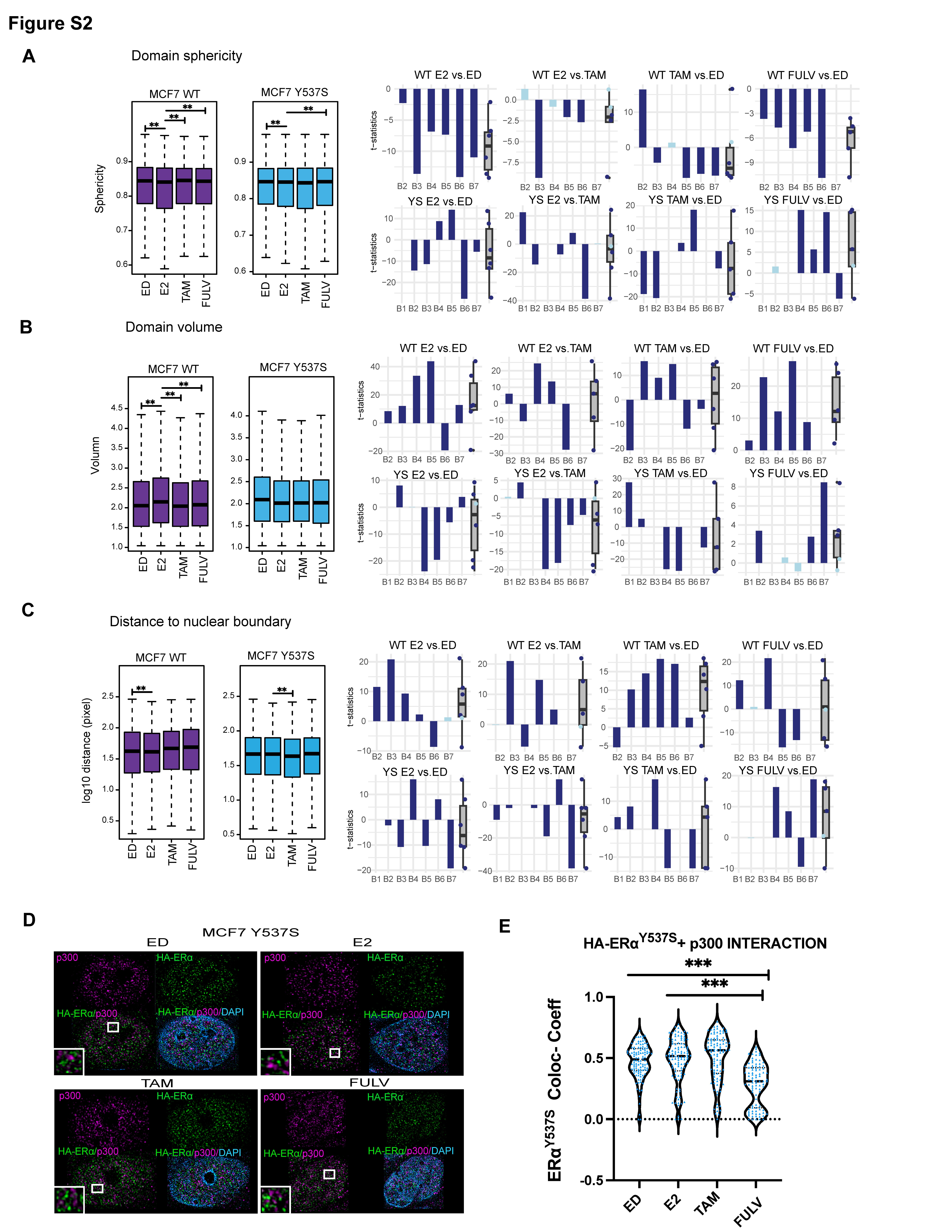

### Supplementary Figure 3

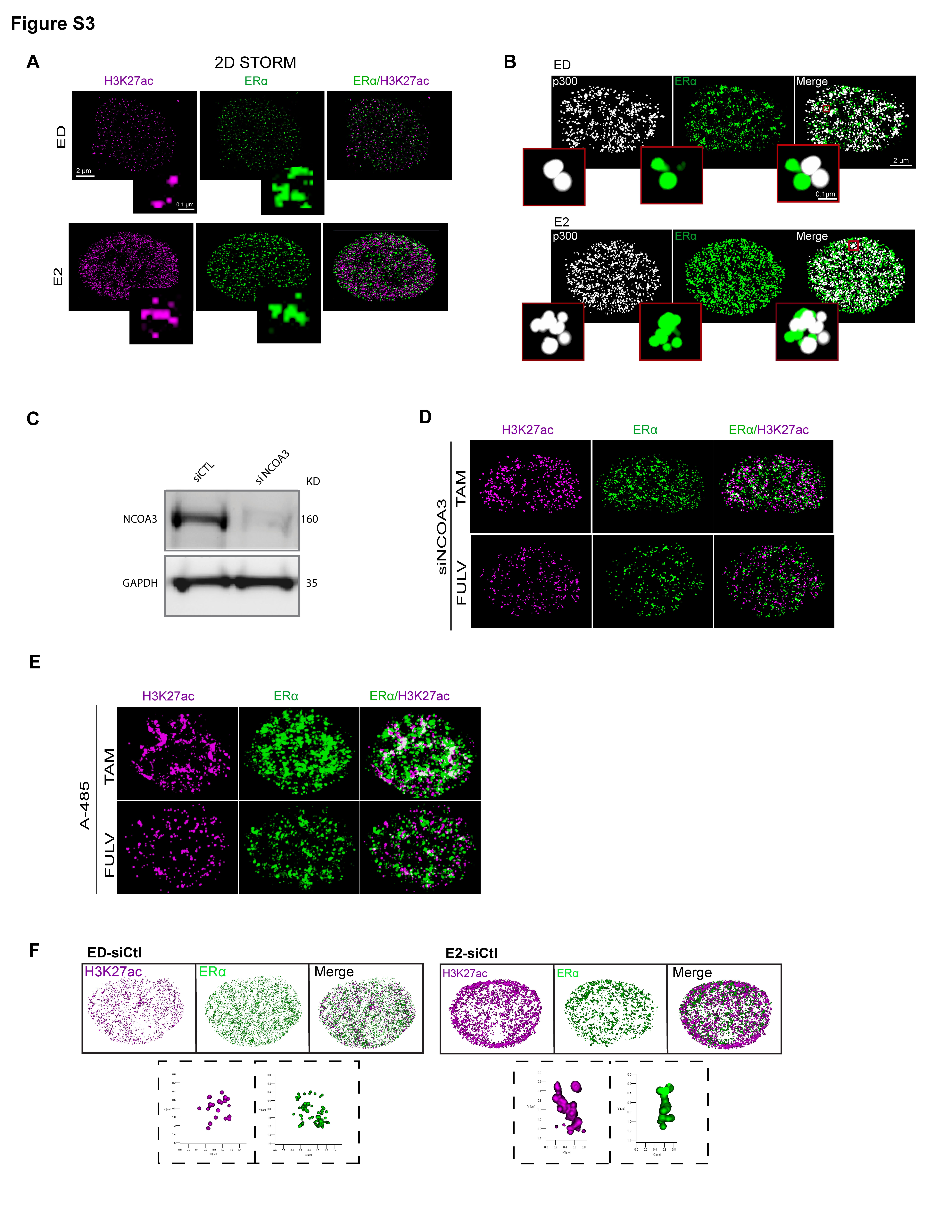

### Supplementary Figure 4

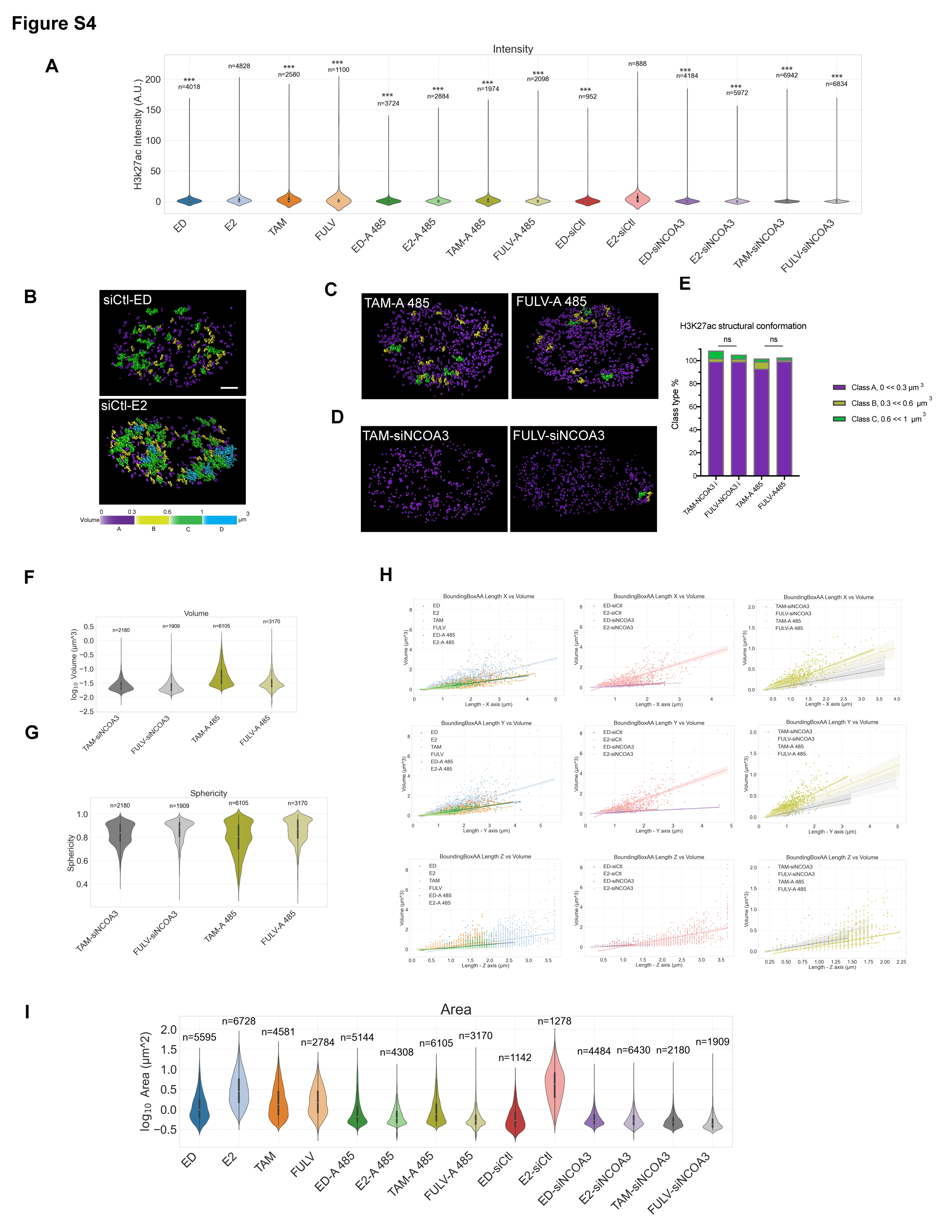

### Supplementary Figure 5

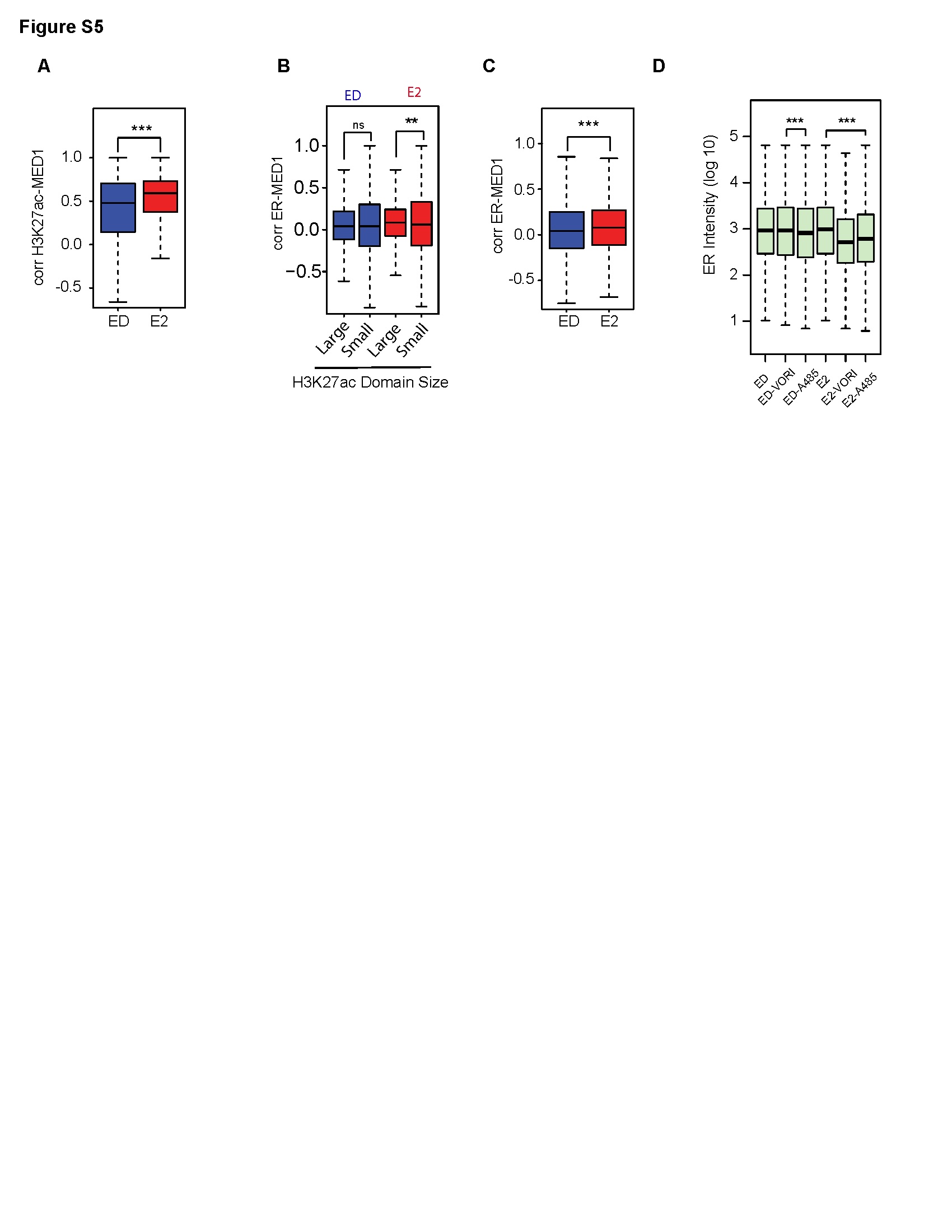

### Supplementary Figure 6

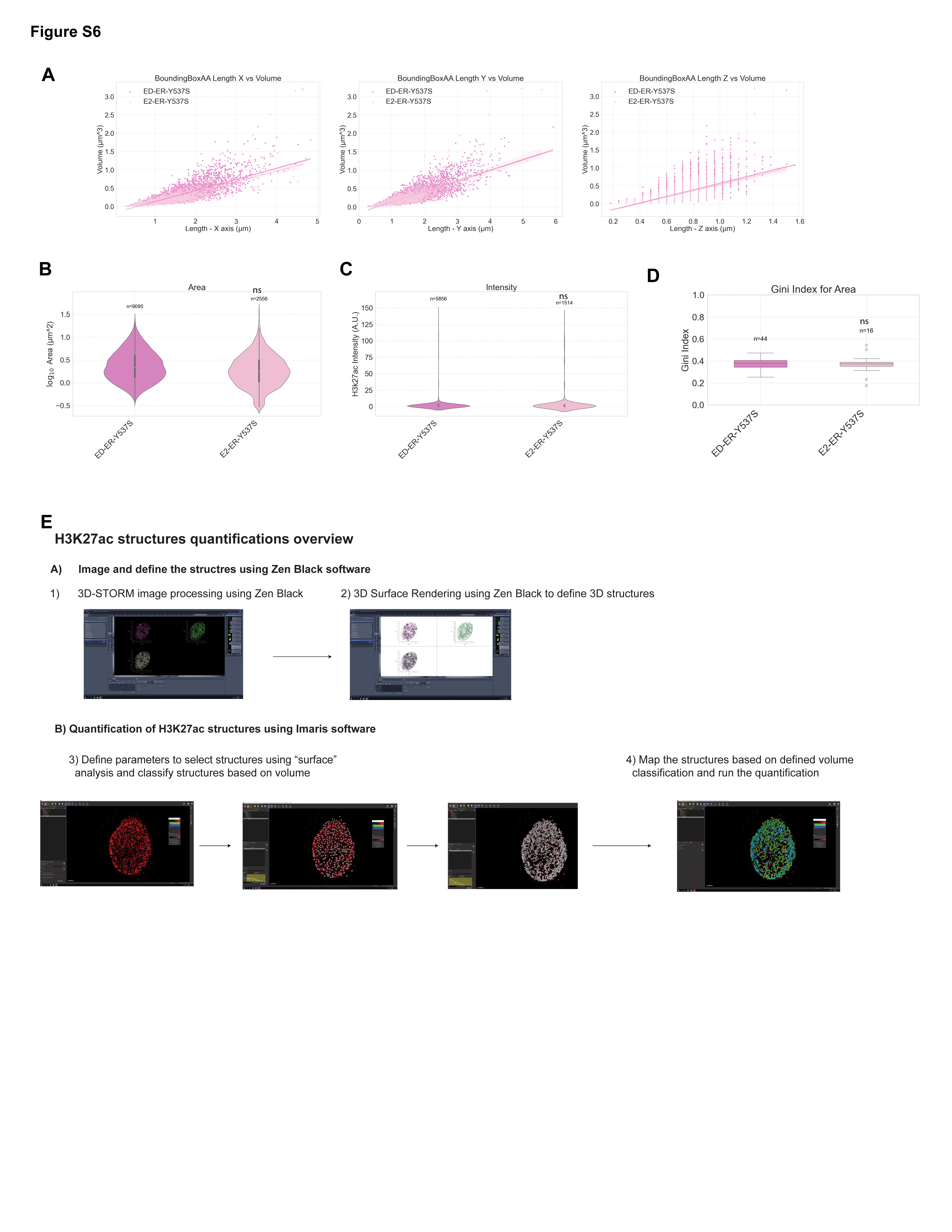
