## Supplementary Information for "Super-resolution microscopy reveals distinct epigenetic states regulated by estrogen receptor activity"

**Materials and Methods**

**Cell Culture**

MCF7 cells were cultured in Dulbecco’s Modified Eagle Medium (DMEM, Gibco, Cat# 11965092) supplemented with 10% fetal bovine serum (FBS, Gibco, Cat# 10437028), 1% penicillin-streptomycin (Gibco, Cat# 15140122), and 1% L-glutamine (Gibco, Cat# 25030081) at 37°C in a humidified 5% CO₂ incubator. For hormone deprivation (HD), cells were cultured in phenol red-free DMEM (Gibco, Cat# 21063029) with 10% charcoal-stripped FBS (Gibco, Cat# 12676029) for 72 hours. Treatments included estradiol (E2, Sigma-Aldrich, Cat# E2758, 10 nM), tamoxifen (Tam, Sigma-Aldrich, Cat# T5648, 100 nM), fulvestrant (Fulv, Sigma-Aldrich, Cat# I4409, 100 nM), and the p300 inhibitor A-485 (MedChemExpress, Cat# HY-100231, 80 nM). Doxycycline (DOX)-inducible ER Y537S mutant MCF7 cells were generated as previously described [46].

**siRNA and Drug Treatments**

siRNA transfection was performed using Lipofectamine RNAiMAX (Invitrogen, Cat# 13778150). Cells were transfected with siRNA targeting NCOA3 (Invitrogen Silencer Select, Cat# s15698) or non-targeting control siRNA (Invitrogen Silencer Select Negative Control No. 1, Cat# 4390843) for 48 hours prior to treatment. For drug treatments, cells were treated with E2, Tam, or Fulv for 45 minutes to 1 hour, or with A-485 for 3 hours prior to E2 stimulation.

**Immunofluorescence (IF)**

Cells were fixed with cold methanol (Fisher Scientific, Cat# A412-4) for 15 minutes at 4°C and blocked with 5% normal donkey serum (Abcam, Cat# ab7475) for 30 minutes. Primary antibodies against ERα (Sigma-Aldrich, Prestige Antibodies, Cat# HPA000449), H3K27ac (Invitrogen, Cat# MA5-23516), and p300 (Invitrogen, Cat# MA5-14894) were applied for 90 minutes at room temperature, followed by fluorophore-conjugated secondary antibodies (Alexa Fluor 488, Invitrogen, Cat# A-21206; Alexa Fluor 647, Invitrogen, Cat# A-31573) for 1 hour. For STORM imaging, cells were maintained in PBS and plated in ibidi 35 mm imaging dishes (ibidi, Cat# 81156). For confocal and SIM imaging, cells were seeded on #1.5 thickness round cover glasses (Fisher Scientific, Cat# 12-545-80), stained with DAPI (Invitrogen, Cat# D1306), and mounted using ProLong Glass Mounting Media (Invitrogen, Cat# P36980).

**Western Blot**

Cells were lysed in RIPA buffer (Thermo Fisher, Cat# 89900), and protein concentrations were determined using a BCA assay (Thermo Fisher, Cat# 23225). Proteins were separated by SDS-PAGE, transferred to PVDF membranes, and probed with antibodies against NCOA3 (Cell Signaling Technology, Cat# 14234) and β-actin (loading control, Cell Signaling Technology, Cat# 4970). Blots were developed using chemiluminescence.

**Imaging**

**Confocal Microscopy**

Confocal images were acquired using a Zeiss LSM 980 AiryScan microscope with a 63x oil immersion objective (NA 1.4). Z-stacks were collected with a step size of 0.15 µm.

**Structured Illumination Microscopy (SIM)**

3D-SIM images were acquired using a Zeiss Elyra PS.1 system equipped with a 63x oil immersion objective (Plan-Apochromat 63x/1.40 Oil DIC, Zeiss, Cat# 420782-9900-000) and dual PCO.edge 4.2 CLHS cameras. Raw images were reconstructed using ZEN Black software, with pixel spacing of 0.06 µm (X/Y) and 0.15 µm (Z-axis). Fluorophores were activated using 405 nm, 488 nm, and 647 nm lasers.

**Stochastic Optical Reconstruction Microscopy (STORM)**

STORM imaging was performed using a Zeiss Elyra PS.1 system equipped with a 63× oil-immersion objective (Plan-Apochromat 63x/1.40 Oil DIC M27, WD=WD=0.19mm), Dual pco.edge 4.2 sCMOScamera. Samples were imaged in dSTORM mode with a freshly prepared imaging buffer containing pyranose oxidase (5 U/mL; Sigma-Aldrich, #P4234), catalase (40 µg/mL; Sigma-Aldrich, #C40), glucose (10% w/v; Sigma-Aldrich, #G8270), and β-mercaptoethanol (100 mM; Sigma-Aldrich, #M6250) to facilitate photoswitching. The buffer replaced PBS immediately prior to imaging.

For each acquisition, 488 nm & 647 nm laser excitation was used, and fluorescence emission was collected through BP 570-620 + LP 655, and BP 420-480 + BP 495-550 nm filters. Images were acquired over 30,000–70,000 frames, with acquisition halted once photon counts dropped to 50% of initial levels to ensure optimal localization precision. Raw data were reconstructed using the ZEN Black software (Zeiss) with default parameters for drift correction and localization analysis.

**Data Processing**

**Balanced Voxel Dimensions Using Z-axis Interpolation**

Raw 3D-SIM images have a pixel spacing of 0.06 µm in the X/Y plane (X/Y-axes) and a spacing of 0.15 µm between adjacent slides (Z-axis). To address the anisotropic voxel dimensions between the X/Y and Z axes, we applied our previously developed interpolation method [31]. Specifically, we interpolated two equidistant slides between each pair of adjacent real Z-slides along the Z-axis. The signal intensity of each interpolated slide, denoted as Is^*Is*^, was estimated as the weighted average of the two neighboring real slides:

$\hat{I_{s}}=\frac{2}{3}I_{s0}+\frac{1}{3}I_{s1}$ (1)

where I_s0_*I_s_*_0_ and I_s1_*I_s_*_1_ represent the signal intensities of the closer and farther real slides, respectively. The weights (2/3 and 1/3) were assigned based on the relative distances of the interpolated slide from the adjacent real slides. The two interpolated slides were positioned at 0.05 µm intervals, equidistant from the real slides. This approach transformed the original 0.15 µm spacing between real Z-slides into a sequence of four slides (two real and two interpolated) with uniform 0.05 µm spacing. Consequently, the interpolation increased the Z-axis resolution, aligning it (0.05 µm) more closely with the X/Y-axis spacing (0.06 µm). The interpolated (enhanced) image data were used for subsequent analysis.

**Detection of Cell Nucleus Regions**

To identify cell nucleus regions in the enhanced 3D-SIM image data, we performed 2D segmentation for each slide independently, rather than applying a 3D approach, due to the anisotropic voxel dimensions. Although Z-axis resolution was improved through interpolation (from 0.15 µm to 0.05 µm), the overall Z-axis length remained significantly shorter than the X/Y dimensions (e.g., 10–20 real slides versus 1500 × 1500 pixels per slide). Thus, a 2D slide-by-slide analysis was chosen to ensure accurate and robust nucleus detection (Figure S1A).

For each slide, we used the DAPI channel signal intensity to detect cell nucleus regions. The DAPI signal intensity was thresholded to generate a binary mask, where pixels with intensities > 500 were classified as “ON” pixels for potential nucleus regions (Figure S1A). Next, we connected the ON pixels in the binary mask as connected components using an 8-connectivity neighborhood structure. The top 15 largest components/regions were selected and processed. For each of these components, we performed morphological operations, including hole filling, dilation (10 iterations), and a second hole filling, to refine the region detection. The refined region was defined as a cell nucleus if it had a pixel size ≥ 100,000 and a sphericity score > 0.4 (sphericity explained below) (Figure S1A).

To detect the boundaries of each cell nucleus, we applied the Sobel operator in the X and Y directions, followed by combined gradient detection. The resulting edge map was binarized to highlight nucleus boundaries, and boundary positions were extracted (Figure S1A).

The 2D cell nucleus regions and boundaries were then stacked across all slides (real and interpolated) to form the final 3D nucleus regions and boundaries for downstream analysis. This approach ensured accurate nucleus detection and boundary identification on a per-slide basis while leveraging the enhanced Z-axis resolution for consistent 3D reconstruction.

**Detection of Chromatin Domains**

We employed a modified version of our previous method [29] to detect 3D H3K27ac-associated chromatin domains within the defined cell nucleus regions. First, we used a data-driven thresholding approach based on H3K27ac signal intensity to identify “ON” pixels. We collected H3K27ac signal intensities from pixels outside the nucleus regions (background distribution; Figure S1B, blue histogram). The top 0.1 percentile value of this background distribution was chosen as the threshold for “ON” pixels, effectively distinguishing high H3K27ac signal regions. This cutoff was then applied to pixels within the nucleus (Figure S1B, red histogram). Pixels with intensities above this threshold were classified as “ON,” indicating active chromatin regions.

Next, we connected the ON pixels as 3D connected components using an 8-connectivity neighborhood structure. We performed morphological operations, including hole filling, dilation (two iterations), and a second hole filling, to refine the components. The resulting regions were filtered by size, retaining those with pixel counts between 10 and 100,000 (the expected range for chromatin domains). This process enabled accurate detection and segmentation of H3K27ac-associated chromatin domains for subsequent feature analysis.

**Channel Alignment**

We observed a systematic shift between the H3K27ac and ER channels (Figure S1C, upper) and designed a correction method to improve alignment. Specifically, we calculated a 2D cross-correlation matrix (Pearson correlation coefficient) between the H3K27ac and ER channel images for each slide. The peak value in this matrix, corresponding to the highest correlation coefficient, identified the optimal relative shift in the X/Y directions. The ER channel was then adjusted to this new position (Figure S1C, lower), and the aligned ER channel was used for subsequent comparisons.

**Feature Collection for H3K27ac Chromatin Domains**

The following features are collected for each of the H3K27ac chromatin domains detected:

1. The correlation coefficient between H3K27ac and ER signal intensities (Figure 1C,D, Figure S1D, E). For each chromatin domain detected, we collect H3K27ac and ER signal intensities for all pixels in the domain and calculate the Pearson correlation coefficient between these two signal vectors. The higher correlation coefficient indicates a better association between H3K27ac occupancy and ER binding in the target domain.
2. The volume of the domain (Figure 1E, Figure S2A), measured as the pixel count of the domain.
3. The sphericity of the domain (Figure 1F, Figure S2A). For each chromatin domain detected, we calculate the 3D sphericity ($\phi$) based on the 3D spatial distribution of the domain pixels with the following formula:

$$\phi=\frac{\pi^{\frac{1}{3}}{(6\times V)}^{2/3}}{S} (2)$$

Where $S$ is the surface area of the domain, and $V$ is the volume of the domain. This metric provides an intuitive measure of domain shape, with higher sphericity values indicating a denser chromatin structure and lower sphericity values for a looser chromatin structure.

4. The distance of the domain to the nearest nuclear boundary (Figure 1G, Figure S2C). Chromatin domains further away from the nuclear boundary indicate a higher possibility of locating within the TAD domain and being more active. Domains closer to the boundary indicates a higher possibility of locating within the lamina domain and being repressive/dense.

For each feature, we compare their values among samples from different conditions (e.g., ED, E2, TAM, and FULV) and genotypes (e.g., wild type and Y537S mutant). We perform this comparison in both batch-combined levels (Figure 1C, Figure 1E-G, and Figure S2A-C left panels) and batch-separated levels (Figure 1D, Figure 1E,F Figure S1D, and Figure S2A-C right panels). We use a one-sided t-test to examine whether the features are higher in E2 than in any other conditions, and the significance is labeled in the figures.

In order to explore the association between domain size and the association between H3K27ac and MED1 signal, we separate H3K27ac domains into small (<=100 pixels) and large (>100 pixels) groups and check the distribution (boxplots) of the Pearson correlation coefficient between H3K27ac and MED1 pixel-level signal within each H3K27ac domain.

**Data Analysis Using Imaris Software**

For volume analysis and 3D rendering, raw images acquired using STORM were processed using Imaris software (version 10, Oxford Instruments). Individual chromatin structures were selected using the surface function, with parameters based on intensity and volume. Structures were categorized into four volume-based classes, and further analyses (e.g., area, intensity) were performed (Fig S6E). Data were exported for statistical analysis and visualization using Python, including Gini index calculations and plot generation.

**Statistical Analysis**

We used one-sided Student T-tests to examine the significance of the difference in 3DSIM data. Significance levels were labeled as: *, p < 0.05; **, p < 0.01; ***, p < 0.001. The T-statistics were also summarized for different batches and displayed as boxplots (Figures 1D, S1D).

Data from STORM processing software were analyzed using Python. Violin plots visualized the distribution of chromatin structure attributes (volume, area, intensity, sphericity), while scatter plots examined relationships between structure length and volume, with linear regression lines. Due to non-Gaussian data distribution (confirmed by Shapiro-Wilk and Kolmogorov-Smirnov tests), non-parametric tests were used. A Kruskal-Wallis ANOVA assessed differences in medians across conditions (e.g., HD, E2, Tam, Fulv, genotypes). Significant results (p < 0.05) were followed by Mann-Whitney U tests with Bonferroni correction for pairwise comparisons. Significance levels were annotated as: p < 0.05 = *, p < 0.01 = **, p < 0.001 = ***.

To evaluate cellular heterogeneity, the Gini index was calculated for each attribute, measuring inequality within individual cells (0 = perfect equality, 1 = total inequality). Gini index distributions across conditions were compared using Kruskal-Wallis and Mann-Whitney tests. Analyses were performed on both batch-combined and batch-separated datasets, with significant differences marked in plots.

**Supplementary data**

**Figure S1**

**(A)** Detection of cell nuclei using DAPI signal (visualized in a single Z stack). Left panel for grayscale representation of DAPI intensity, mid panel for identification of the DAPI ON/OFF pixels (white/black), and right panel for identified cell nucleus boundaries, and H3K27ac signal histogram with cutoff (dashed line).
**(B)** Pixel-intensity distribution of H3K27ac signals in nuclear (red) versus non-nuclear (blue) regions from a representative 3D-SIM image. Dashed line marks the 99.9th percentile cutoff for H3K27ac-positive pixels.
**(C)** 2D cross-correlation coefficient between channel 1 signal (H3K27ac) and channel 2 signal (ER) before (upper panel) and after (lower panel) correction for a 3D-SIM sample. The X and Y axes represent the relative shift size of channel 2 (given the fixed channel 1) on the X and Y axes, respectively. The lighter colors represent a higher correlation coefficient, and the darker colors represent a lower correlation coefficient.
**(D/E)** t-statistic from each of the six batches (six bars) in ER-WT (D) and ER-Y537S mutant cells (E)The t-statistic quantifies the difference in correlation between H3K27ac and ER signal intensity in E2 compared to ED, considering all H3K27ac domains detected in the samples from each corresponding batch. A boxplot summarizing the six bars was also attached. Different panels represented different comparisons between conditions. Bars in dark blue and light blue colors represent P < 0.05 and non-significant (by t-test), respectively. Positive and negative scores indicate higher and lower correlation coefficients in E2-treated cells, respectively.
**(F)** SIM images of MCF7 YS cells under ED, E2, Tam, and Fulv conditions, showing ERα-H3K27ac proximity.

**Figure S2**

**(A)** Boxplots on the left showing the H3K27ac domain sphericity across conditions in wild-type and Y537S mutant MCF7 cells, with t-test showing the lower sphericity in E2. P-values from t-test were provided for E2 compared to other conditions. Barplots/boxplots on the right showing the t-statistic testing the difference of H3K27ac domain sphericity in MCF7 cells across conditions and genotypes. Positive and negative scores indicate higher and lower domains sphericity in E2-treated cells, respectively.
**(B, C)** Similar to (A) but for domain volume (B) and distance to the nuclear boundary (C) metrics.

**D)** SIM images of MCF7 YS cells under ED, E2, Tam, and Fulv conditions, showing p300 and H3K27ac proximity.

**E)** Violin plots of colocalization coefficients between p300 and ERα in wild-type and Y537S mutant MCF7 cells under different conditions.

**Figure S3**

**(A)** 2D-STORM images of H3K27ac in ED and E2-treated MCF7 cells, highlighting resolution superiority over SIM.

**(B)** 3D-STORM images of MCF7 cells with/without E2, probed for p300 (gray) and ERα (green), showing p300-ERα correlation in E2-treated cells

**(C)** Western blot showing NCOA3 knockdown efficiency.
**(D)** STORM images of MCF7 cells treated with Tam and Fulv upon NCOA3 knockdownNCOA3 siRNA.

**(E)** Rendered STORM images of control siRNA cells in ED and E2 conditions.

**(F)** STORM images of MCF7 cells treated with Tam and Fulv in combination with A485 (80 nM).

**Figure S4**

**(A)** Violin plot showing changes in global H3K27ac levels across conditions.

**(B)** Structural maps of control siRNA cells (±E2), classified by volume (yellow/green/blue: high; purple: low)

**(C-E)** Structural maps (C, D) and class distributions (E) for each conditions as indicated.

**(F-G)** Quantitative analyses of volume (F), sphericity (G) for each conditions as indicated.
**(H)** Volume quantification across X, Y, Z axes for all conditions as indicated.
**(I)** Area quantifications for H3K27ac structures for each conditions as indicated.

**Figure S5**

**(A)** Boxplots of pixel-level correlation coefficients between MED1 and total H3K27ac domains, regardless of size.
**(B)** Boxplots of pixel-level correlation coefficients between MED1 and ERα, categorized by large versus small H3K27ac structures.
**(C)** Boxplots of pixel-level correlation coefficients between MED1 and total ERα, regardless of domain size.
**(D)** Quantification of ERα signal intensity under each condition as indicated.

**Figure S6**

**(A, B)** Volume quantification across X, Y, Z axes (A) and projected areas (B) for Y537S cells with/without E2.

**(C)** Violin plot showing no change in global H3K27ac levels in ED vs. E2 conditions.

**(D)** Gini index for H3K27ac volume heterogeneity in ED vs. E2 conditions.

**(E)** Schematic of 3D reconstruction: STORM rendering (Zen Black) → volume detection → Imaris classification.

**Movie 1 &2**

**1.** Rendered structure of a 3D-STORM image of an MCF-7 cell treated with 10 nM E2 for 1 hour following 3 days of estrogen deprivation (ED). ERα is depicted in green, and H3K27ac is shown in magenta. **2.** Magnified view highlighting the spatial organization of ERα and H3K27ac within the same cell.

**Movie 3 &4**

**3.** Rendered structure of a 3D-STORM image of an MCF-7 cell after 3 days of ED. ERα is depicted in green, and H3K27ac is shown in magenta. **4.** Magnified view highlighting the spatial organization of ERα and H3K27ac within the same cell.

**Movie 5 &6**

**5.** Rendered structure of a 3D-STORM image of an MCF-7 cell treated with 100 nM Tam for 1 hour following 3 days of ED. ERα is depicted in green, and H3K27ac is shown in magenta. **6.** Magnified view highlighting the spatial organization of ERα and H3K27ac within the same cell.

**Movie 7 &8**

**7.** Rendered structure of a 3D-STORM image of an MCF-7 cell treated with 100 nM Fulv for 1 hour following 3 days of ED. ERα is depicted in green, and H3K27ac is shown in magenta. **8.** Magnified view highlighting the spatial organization of ERα and H3K27ac within the same cell.

**Movie 9 &10**

**9.** Rendered structure of a 3D-STORM image of an MCF-7 cell treated with 10 nM E2 for 1 hour following 48 hours of NCO3 siRNA and 3 days of ED. ERα is depicted in green, and H3K27ac is shown in magenta. **10.** Magnified view highlighting the spatial organization of ERα and H3K27ac within the same cell.

**Movie 11 &12**

**11.** Rendered structure of a 3D-STORM image of an MCF-7 cell treated with 10 nM E2 for 1 hour following 3 days of ED and 3 hours of A-485 (80 nM). ERα is depicted in green, and H3K27ac is shown in magenta. **12.** Magnified view highlighting the spatial organization of ERα and H3K27ac within the same cell.

**Table S1: Cell number, SIM, H3K27ac analysis**

| **Cell line** | **Treatment** | **H3K27ac domain Number** | **Cell Number** |
| --- | --- | --- | --- |
| **MCF7 WT** | ED | 129,059 | 403 |
|  | E2 | 170,496 | 451 |
|  | TAM | 119,761 | 351 |
|  | FULV | 130,983 | 387 |
| **MCF7 Y537S** | ED | 180,382 | 616 |
|  | E2 | 196,626 | 545 |
|  | TAM | 142,453 | 496 |
|  | FULV | 110,013 | 471 |

**Table S2: Cell number, SIM, p300 analysis**

| **Cell line** | **Treatment** | **Cell Number** |
| --- | --- | --- |
| **MCF7 WT** | ED | 210 |
|  | E2 | 205 |
|  | TAM | 144 |
|  | FULV | 155 |
| **MCF7 Y537S** | ED | 112 |
|  | E2 | 110 |
|  | TAM | 105 |
|  | FULV | 93 |

**Table S3: Cell number, STORM, H3K27ac analysis**

| **Cell line** | **Treatment** | **H3K27ac domain Number** | **Cell Number** |
| --- | --- | --- | --- |
| MCF7 WT | ED | 5595 | 26 |
|  | E2 | 628 | 32 |
|  | TAM | 4581 | 21 |
|  | FULV | 2784 | 15 |
|  | ED -CTL siRNA | 1142 | 15 |
|  | E2- CTL siRNA | 1278 | 15 |
|  | ED -NCOA3 siRNA | 4484 | 21 |
|  | E2-NCOA3 siRNA | 6430 | 32 |
|  | TAM -NCOA3 siRNA | 2180 | 27 |
|  | Fulv-NCOA3 siRNA | 1909 | 28 |
|  | ED -A-485 | 5144 | 25 |
|  | E2-A-485 | 4308 | 24 |
|  | TAM -A-485 | 6105 | 18 |
|  | Fulv-A-485 | 3170 | 15 |
| **MCF7 Y537S** | ED | 9095 | 44 |
|  | E2 | 2556 | 16 |

**Table S4: Cell number, SIM, MED1 analysis**

| **Cell line** | **Treatment** | **H3K27ac domain Number** | **Cell Number** |
| --- | --- | --- | --- |
| **MCF7 WT** | ED | 17313 | 66 |
|  | E2 | 54273 | 146 |

**Table S5: Cell number, SIM, H3K27ac analysis**

| **Cell line** | **Treatment** | **H3K27ac domain Number** | **Cell Number** |
| --- | --- | --- | --- |
| **MCF7 WT** | ED | 17313 | 66 |
|  | E2 | 54273 | 146 |
|  | ED-A485 | 25254 | 52 |
|  | E2-485 | 63090 | 252 |
|  | ED- Vorinostat | 15363 | 73 |
|  | E2- Vorinostat | 26165 | 193 |
